# An inhibitory plasticity mechanism for world structure inference by hippocampal replay

**DOI:** 10.1101/2022.11.02.514897

**Authors:** Zhenrui Liao, Darian Hadjiabadi, Satoshi Terada, Ivan Soltesz, Attila Losonczy

## Abstract

Memory consolidation assimilates recent experiences into long-term memory. This process requires the replay of learned sequences, though the content of these sequences remains controversial. Recent work has shown that the statistics of replay deviate from those of experience: stimuli which are experientially salient may be either selected or suppressed. We find that this phenomenon can be explained parsimoniously and biologically plausibly by a Hebbian spike time-dependent plasticity rule at inhibitory synapses. Using spiking networks at three levels of abstraction–leaky integrate-and-fire, biophysically detailed, and abstract binary–we show that this rule enables efficient inference of a model of the structure of the world. We present analytical results that these replayed sequences converge to ground truth under a mathematical model of replay. Finally, we make specific predictions about the consequences of intact and perturbed inhibitory dynamics for network dynamics and cognition. Our work outlines a potential direct link between the synaptic and cognitive levels of memory consolidation, with implications for both normal learning and neurological disease.

## Introduction

Humans and other animals have the remarkable ability to learn about the world around them from incomplete observations containing a mix of structural features of the environment and distractions that do not generalize. The ability to learn arbitrary sequences and relational maps, and deploy them to guide behavior, is the defining computational function of the hippocampus (Kumaran & Maguire 2005, Eichenbaum 2017, Eichenbaum & Cohen 2014, Whittington et al. 2020, Behrens et al. 2018). A subset of hippocampal neurons, termed “place cells”, exhibit selective tuning to specific locations in space: as an animal explores space, the sequence of place cell firing reflects the animal’s path (O’Keefe & Dostrovsky 1971). The hippocampus later reinstates previously learned sequences in temporally compressed “replay” epochs inside high-frequency oscillatory events known as sharp-wave ripples (SPW-Rs) while the animal is at rest (Buzsaki 2015, Dragoi & Buzsáki 2006, Foster & Wilson 2006, Diba & Buzsaki 2007, Wilson & McNaughton 1994, Lee & Wilson 2002). This replay is required for the long-term consolidation of memories (Girardeau et al. 2009, Ego-Stengel & Wilson 2010, Jadhav et al. 2012, Fernández-Ruiz et al. 2019).

The CA3 region of the hippocampus, characterized by strong recurrent connectivity (Amaral & Witter 1989, Káli & Dayan 2000, Guzman et al. 2016, Rebola et al. 2017), is thought to be the source of SPW-Rs and the driver of replay throughout the hippocampus (Buzsaki 2015, Davoudi & Foster 2019, Csicsvari et al. 1999). Experimental work has recently demonstrated that synaptic weight modification of CA3 recurrent excitatory connections occurs by a symmetric spike-timing dependent plasticity (sSTDP) rule (Mishra et al. 2016), providing a plausible explanation for temporally reversed (“backwards”) replay sequences (Pfeiffer 2020, Foster & Wilson 2006). At the same time, modeling work has demonstrated the feasibility of obtaining spontaneous replay in biologically realistic spiking network models (Nicola & Clopath 2019, Ecker et al. 2022, Milstein et al. 2022). As a densely connected recurrent network with writeable weights, CA3 is a natural system in which to study the content and function of replay with respect to learning from a computational perspective. Nonetheless, CA3 has received much less study than CA1.

Comparatively little is know about inhibitory dynamics in CA3, although inhibition is known to play a major role in SPW-R generation (English et al. 2014, Stark et al. 2014). Under the classic theoretical view of learning, Hebbian plasticity remodels excitatory-excitatory synapses against a relatively static inhibitory background (Hebb 1949, Marr et al. 1991). Plasticity at the GABAergic synapse has traditionally been discounted, under the presumption that a constant inhibitory background was necessary for learning; however, recent experimental and theoretical work has suggested a role for inhibitory plasticity in shaping network-level computations (Chiu et al. 2019, Kullmann & Lamsa 2007, Yap et al. 2021, Udakis et al. 2020, Schulz et al. 2021, Rubin et al. 2017, Vogels & Abbott 2009, Sadeh & Clopath 2021, Loisy et al. 2022).

At the same time, while replay has been characterized at the macroscopic level through electrical or optogenetic manipulations (Gridchyn et al. 2020, Girardeau et al. 2009, Ego-Stengel & Wilson 2010, Jadhav et al. 2012, Fernández-Ruiz et al. 2019) as well as the microscopic level through single-neuron recordings (Valero et al. 2017, English et al. 2014, Maier et al. 2011, Hulse et al. 2016), there is currently no consensus on how replay events emerge spontaneously from a recurrently connected network, or how they aid in memory consolidation. The question of whether predictive representations or specific past experiences are reinstated in replay has been the subject of significant debate (Gillespie et al. 2021). The former model predicts that imminent decisions ought to modulate replay content, while the latter model predicts that replay events should faithfully reactivate sequences representing individual experiences of the environment, even when not immediately behaviorally relevant. However, animals exploring real environments simultaneously experience both consistent features of the world and one-time distractors which do not generalize: thus, memory consolidation is not solely a task of encoding past experience, but also of constructing a model of the world which makes judgments about what is relevant and what is not. Recent work examined SPW-R recruitment of CA3 units in a task combining stochastic, experience-specific stimuli as well as conserved features of the world (Terada et al. 2022), finding that, contrary to the prediction of the specific-experience model of replay, even when salient random stimuli dominate the animal’s specific experiences, representations of those experience-specific stimuli are not only not acutely reactivated but actively and durably filtered out from sharp-wave ripples. Similar work has shown in CA1 that although an animal’s behavior may be strongly biased by reward occupancy, replay content prioritizes unrewarded and less-occupied locations in order to consolidate an unbiased representation of space (Grosmark et al. 2021).

In this work, we investigate a third option in the debate between replay of predictive representations vs specific past experiences: the replay of “world structure”, i.e., the latent statistical relationships between conjunctively coded features of experience. We argue that replay neither reprises specific past experiences, nor solely generates concrete predictions or plans of future behavior, but rather returns samples from the animal’s working model of the deeper structure of its world. This conceptual model synthesizes elements of both of these theories with prior work on model-based replay to explain how replay can be useful for guiding future behavior, even though representations from distal past experience which are not imminently behaviorally relevant are sometimes replayed (McClelland et al. 1995, Pezzulo et al. 2014, Foster 2017). In support of this conceptual framework, we present three levels of modeling—spiking network, biophysical, and abstract normative—reproducing existing experimental findings and demonstrating consolidation of world structure in recurrently connected models of CA3. Central to all three approaches is our major prediction, the use of the spike time-dependent LTP rule to remodel inhibitory synapses, allowing the statistical structure of the world to be learned dynamically from experience. To our knowledge, this is the first work positing a computational role for the long-term potentiation of inhibitory synapses in the hippocampus.

## Results

### Network model of CA3 replay

We first built a large, randomly recurrently connected CA3 spiking network capable of internally generating spontaneous replay (Ecker et al. 2022). We simulated an experiment in which the CA3 network was exposed to both spatial input and random distractor stimuli in an online phase during which synaptic weights were learned by symmetric STDP (sSTDP). Then, in a subsequent offline phase, learned patterns were read out from spontaneous replay (Fig 1a, see (Terada et al. 2022)). Our network consisted of 8000 leaky integrate-and-fire (LIF) pyramidal cells (PCs) and 150 interneurons (INs) in BRIAN2 (Stimberg et al. 2019) with spike-rate adaptation (Gerstner et al. 2014, Ecker et al. 2022), with 50% of the cells assigned to receive place input and a (non-mutually exclusive) 10% receiving stochastic distractor cue input (Terada et al. 2022) (Fig 1b). These neurons were randomly recurrently connected, with a connection probability of 10% between PCs as well as from PCs onto INs, and a connection probability of 25% between INs as well as from INs onto PCs (see **Methods**).

**Figure 1.**
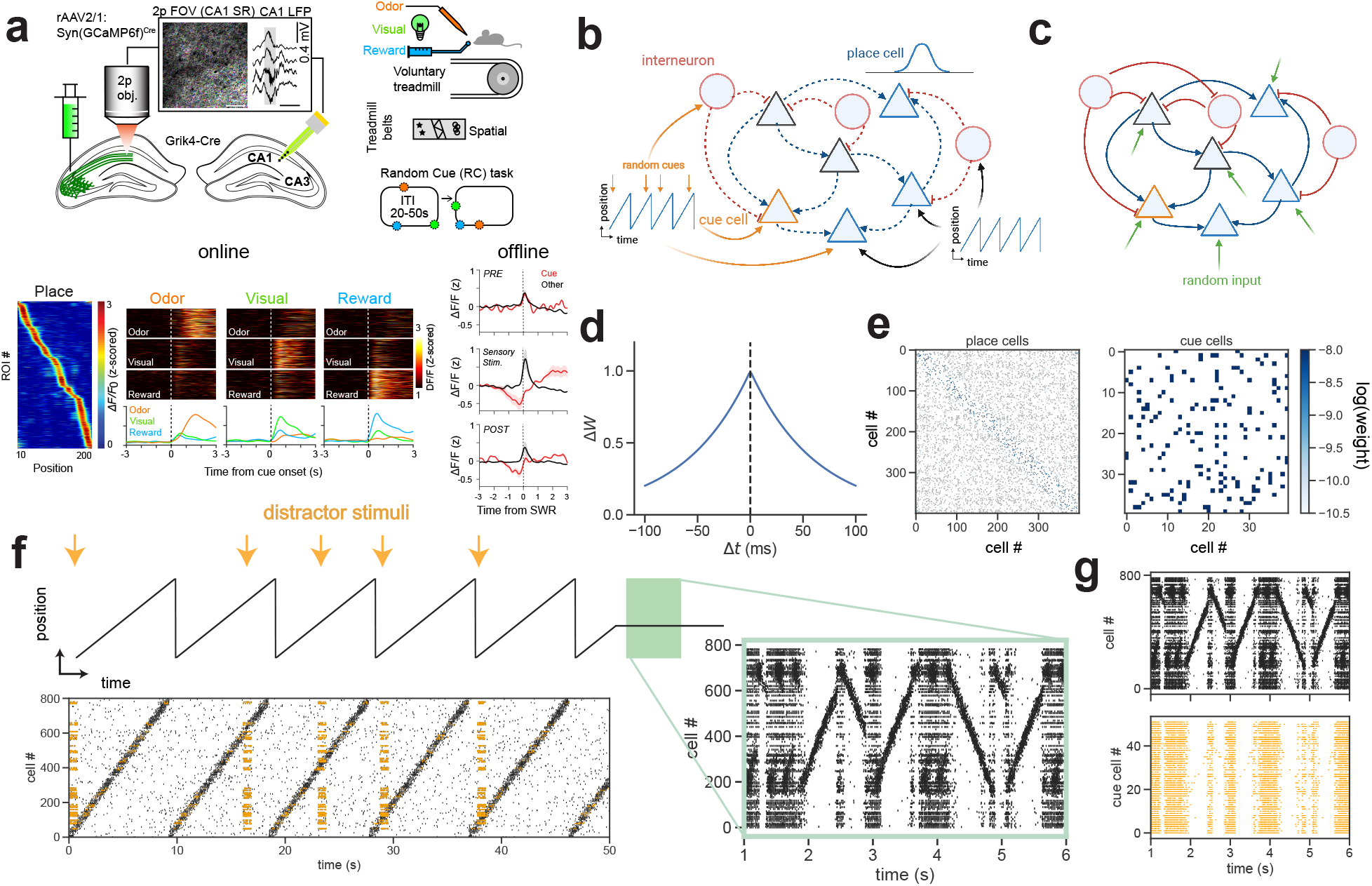
Suppression of distractor stimuli in a random network model of CA3 replay. **a**. Experimental paradigm and results adapted from Terada et al. (2022) (reprinted with permission). Calcium activity of CA3 Schaffer collateral axons was imaged with simultaneous contralateral local field potential (LFP) recordings to detect sharp-wave ripples (SPW-Rs, upper left). Random cue-spatial task (RC-S): animals ran along a circular spatial belt, while three distractor sensory cues (odor, visual, and reward) were presented at pseudorandom locations to the mouse on every lap (upper right). CA3 units robustly represent both space (bottom left) and random cues (bottom center) in the online period. Bottom right: prior to the experiment, space and cue cells are indistinguishable in their SPW-R recruitment. In immobility epochs during the experiment, cue cells are suppressed from SPW-Rs while place cells are recruited, an effect which persists in an immobility recording after the experiment. **b**. Network architecture during learning. Network consists of 8000 PCs (triangles) and 150 INs (circles) randomly recurrently connected. PCs and INs receive random cue and place input. Dashed lines indicate plastic synapses. **c**. Network during replay. Network now only receives random low-level spiking input (green arrows). **d**. Symmetric STDP kernel used for training the network: weight change Δ*W* as a function of temporal difference Δ*t* between pre- and postsynaptic spikes **e**. Learned synaptic weight matrices from a random sample of 400 place cells (left, sorted by location of peak tuning) and 50 cue cells (right, arbitrarily sorted), log scale. Place cells form strong weights to other place cells with peaks shortly before and after their own peak and weak weights with all other cells, while cue cells form strong weights with each other. **f**. Simulation of task in 1a (last 50 seconds of online learning and 10 seconds offline, slice of 10% of the PC network). Top: Simulated position and distractor inputs. Bottom: Raster of network activity during learning, sorted by cells’ tuning curve peak. Cue-responsive cells highlighted in yellow. Right inset: Spontaneous replay emerges in offline period. **g**. Neurons tuned to stochastic distractors (bottom) are active offline, but suppressed during replay.

The simulated experiment consisted of 40 laps through a 3 m long environment at constant velocity. Untuned cells fired at a baseline rate of 0.1 Hz (sampled from a homogeneous Poisson process) throughout the belt. The spike trains of place cells were sampled from an inhomogeneous Poisson process with a position-dependent Poisson rate *λ*(*x*) (maximum 20 Hz) chosen as a circular Gaussian representing the cell’s tuning curve. The spike trains of distractor-tuned cells were sampled in the same way, with all distractor cells sharing a distractor tuning curve, and the distractor appearing stochastically at a randomized location in each lap (Fig 1f top, yellow arrows). Like excitatory neurons, inhibitory neurons in our network may be tuned to place, cue, both, or neither, as experimentally reported (Geiller et al. 2020, Wiebe & Stäubli 2001). During this exploration phase, the weights of this network were trained using sSTDP (Fig 1d, 1f). Example slices from the trained weight matrices are shown in Fig 1e. A diagonal band of strong weight emerges in the place cell subnetwork (sorted by tuning curve peak), indicating that synapses between place cells with adjacent peaks develop strong connections in proportion to the distance between their tuning peaks. Meanwhile, the distractor cell subnetwork becomes strongly recurrently coupled, as distractor cells are coactive with each other (Fig 1e, right).

To simulate the offline state (Fig 1f, green highlight and inset), PCs were driven with random Poisson spiking input irrespective of their distractor-, place-, or untuned status, simulating unstructured input from mossy fibers (Fig 1c). We find that place sequences encountered during learning are spontaneously replayed in the forward and backward directions offline in our network (Fig 1f). Further, we find that the distractor-tuned neurons, despite being generally active in the offline period, are selectively suppressed during spontaneous ripples (Fig 1g). We found that spontaneous replay in the network was robust to variations in the exact *I→E* plasticity rule used, so long as the plasticity rule was a Hebbian LTP rule operating on a timescale of roughly 3 ms (Fig S1, S2).

In summary, we have shown that replay spontaneously emerges in a randomly recurrently connected network trained on a simulated random cue-spatial (RC-S) task using sSTDP of excitatory and inhibitory synapses. We have further shown that even though this replay is completely spontaneous and internal to the network, with no explicit externally applied “replay signal”, and the learning rule is strictly local to individual synapses, the network self-organizes distributed and specific suppression of cue cells during replay epochs.

### Biophysically detailed model of CA3 replay

We next asked whether sSTDP of recurrent *E→E* and *I→E* synapses is sufficient to suppress distractor inputs during offline replay in a biophysical model of the hippocampal CA3 network. We modeled CA3 using 260 multicomparmental and biophysically-detailed pyramidal neurons (Yu et al. 2020) and 30 interneurons with biophysically realistic electrophysiological properties (Bezaire et al. 2016, Bezaire & Soltesz 2013) (see **Methods**). During online learning, PCs and INs receive input structured into distinct dendritic input streams, consistent with in vivo data (Losonczy et al. 2008, Makara & Magee 2013, Druckmann et al. 2014, Bezaire & Soltesz 2013, Adoff et al. 2021).

Specifically, 80% of PCs receive spatially structured place and grid input via a mossy fiber and medial entorhinal cortical (MEC) pathway, respectively (Knierim 2015, Moser et al. 2008, Knierim et al. 2014). The remaining 20% of PCs receive a distractor input via a lateral entorhinal cortical (LEC) pathway (Knierim et al. 2014) (Fig 2a). As in Figure 1, the PCs receiving distractor input (i.e., cue cells) are driven at a randomly selected location during each lap of the linear virtual track. After online learning with the sSTDP rule, place cell firing increased on average to physiologically relevant firing rates (0.264 Hz to 1.087 Hz) and furthermore robustly tiled the track (Fig 2b) with little to no out of field firing (Fig 2c).

**Figure 2.**
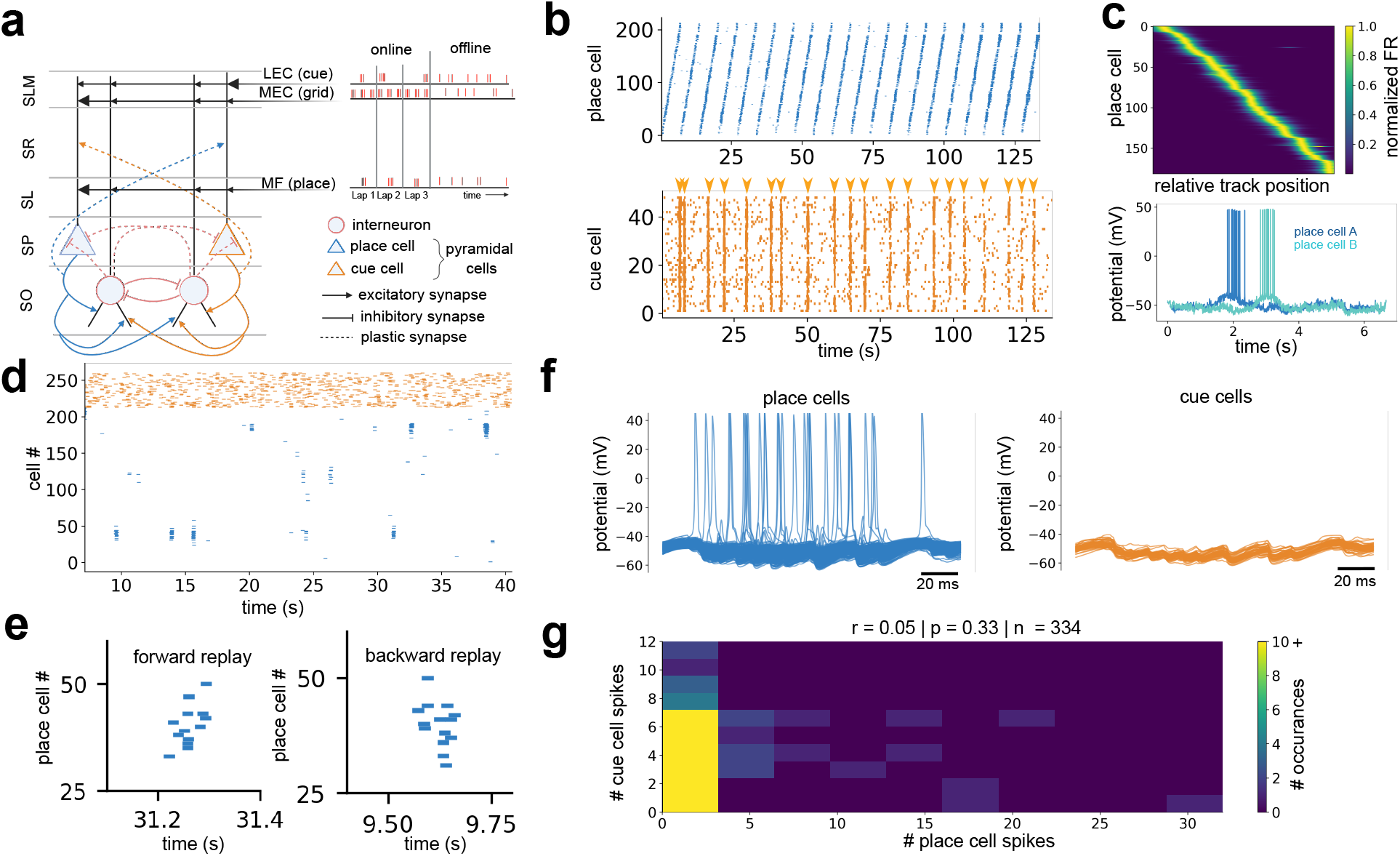
Distractor suppression in a biophysically detailed model of CA3 replay. **a**. Schematic of the biophysical CA3 model. The model consists of excitatory PCs and perisomatic-targeting INs wired in accordance with experimental data. During online learning, the model is driven by multiple sources of structured external input – mossy fibers: place input; medial entorhinal cortex (MEC): grid input; lateral entorhinal cortex (LEC): cue input. During offline replay, the CA3 circuit is driven by random noise. SLM = stratum lacunosum moleculare; SR = stratum radiatum; SL = stratum lucidum; SP = stratum pyramidale; SO = stratum oriens. **b**. Spike raster of place cell (blue) and cue cell (orange) activity during online training during which STDP on *E → E* and *I →E* synapses is active. (left) At the start of training, the place cell ensemble poorly tiles the track and exhibit significant out of field firing. (right) After multiple laps, the place cells effectively tile the linear track and exhibits little to no out of field firing. Orange arrow heads represent instances where cues input is provided to the network. **c**. Top: Place cells are spatially tuned to distinct regions of the linear track after online learning. Bottom:^··^ Somatic voltage potential for two example place cells showing robust in-field firing. **d**. (top) Spike raster of place cells (blue, organized based on peak firing location on linear track) and cue cells (orange, no specific organization) during the offline state. Epochs of high place cell activity show place cells with adjacent tuning peaks firing in rapid succession, consistent with a memory replay event. **e**. Examples of individual place cells spikes during a forward replay (left) and backward replay (right) event organized by peak firing location similar to Fig 2d. Both forward and backward replay event traverse approximately 10% of the length of the linear track and occur on a timescale of ∼100 ms. **f**. Example model replay event during the offline state shows place cells (blue) are active whereas cue cells (orange) are suppressed. **g**. Cue cells and place cells are effectively decoupled during the offline state (Pearson’s *r* = 0.05, *p* = 0.33, *n* = 334).

To investigate the offline dynamics of cue cells and place cells, we simulated one ‘online’ lap during which the network received structured input followed by an ‘offline’ state in which the network received only unstructured random inputs (Fig 2a). During this offline state, epochs of high place cell activity spontaneously emerged (Fig 2d). Sequences of place cells with adjacent tuning peaks along the linear track (Fig 2c) were reactivated during these spontaneous chirps (Fig 2d) in both the forward and backward directions (Fig 2e), suggesting the emergence of spontaneous memory replay. During these replay-like events, cue cells were widely suppressed (Fig 2f), and overall place and cue cell firing was not significantly coupled (Pearson’s *r* = 0.05, *p* = 0.33, *n* = 334) over the course of 30 seconds of the simulated offline state (Fig 2g). Thus, using a data-driven biophysical model of hippocampal CA3 trained with *E→E* and *I→E* sSTDP, we found that, consistent with experimental data, memory replay reinstates representations of the structure of the world while suppressing distractor representations.

### Perturbations and predictions from the biophysical network model

Our central synaptic-level prediction is the existence of a Hebbian LTP rule of GABAergic synapses in the hippocampus which is required for the network-level phenomenon of replay and memory consolidation from noisy observations. Here, we performed a series of perturbation simulations to test the hypothesis that inhibitory plasticity is necessary for cue cell suppression and, more generally, world structure learning. Furthermore, the simulations performed here broaden the main findings and generate additional experimentally verifiable predictions that give insight on neural communication across multiple spatial scales.

We first observed that the median inhibitory weight onto cue cells was 38.9% greater than onto place cells (median *w*_*I→cue*_: 0.0025 uS; median *w*_*I→place*_: 0.0018 uS; *p* = 7.4 *×* 10^−66^, one-sided Mann Whitney U-test) (Fig 3a). Therefore, we performed simulated dual-patch clamp recording of IN-PC pairs to measure inhibitory post-synaptic currents (IPSCs) onto PCs and identify the cellular mechanism that enables cue cells to be suppressed during SPW-Rs. PCs were held in voltage clamp mode with holding voltages ranging from −65 mV to −45 mV in +5 mV increments. We observed that IPSCs onto cue cells (*n* = 960 measured pairs) were significantly greater (one-sided Mann Whitney U-test, *p <* 0.001 for all holding voltages) than IPSCs onto place cells (*n* = 4240 measured pairs) (Fig 3b). The IPSCs onto cue and place cells was smallest at holding potentials closer to the chloride reversal potential (*E*_*rev,Cl*_ = −65 mV, place: 0.31 *±* 0.19 pA; -65 mv, cue: 0.36 *±* 0.07 pA) compared to more depolarized holding potentials (−45 mV, place: 0.94 *±* 0.55 pA; −45 mV, cue: 1.07 *±* 0.21 pA). These simulations predict that sSTPD of GABAergic synapses generates powerful feedforward inhibition onto cue cells (Fig 3c) that may be critical for suppressing cue cells during periods of high place cell activity, such as SPW-Rs. The validity of this inhibitory feedforward motif can be directly tested in future experiments by measuring IPSCs onto place and cue cells from different classes of INs *in vivo*.

**Figure 3.**
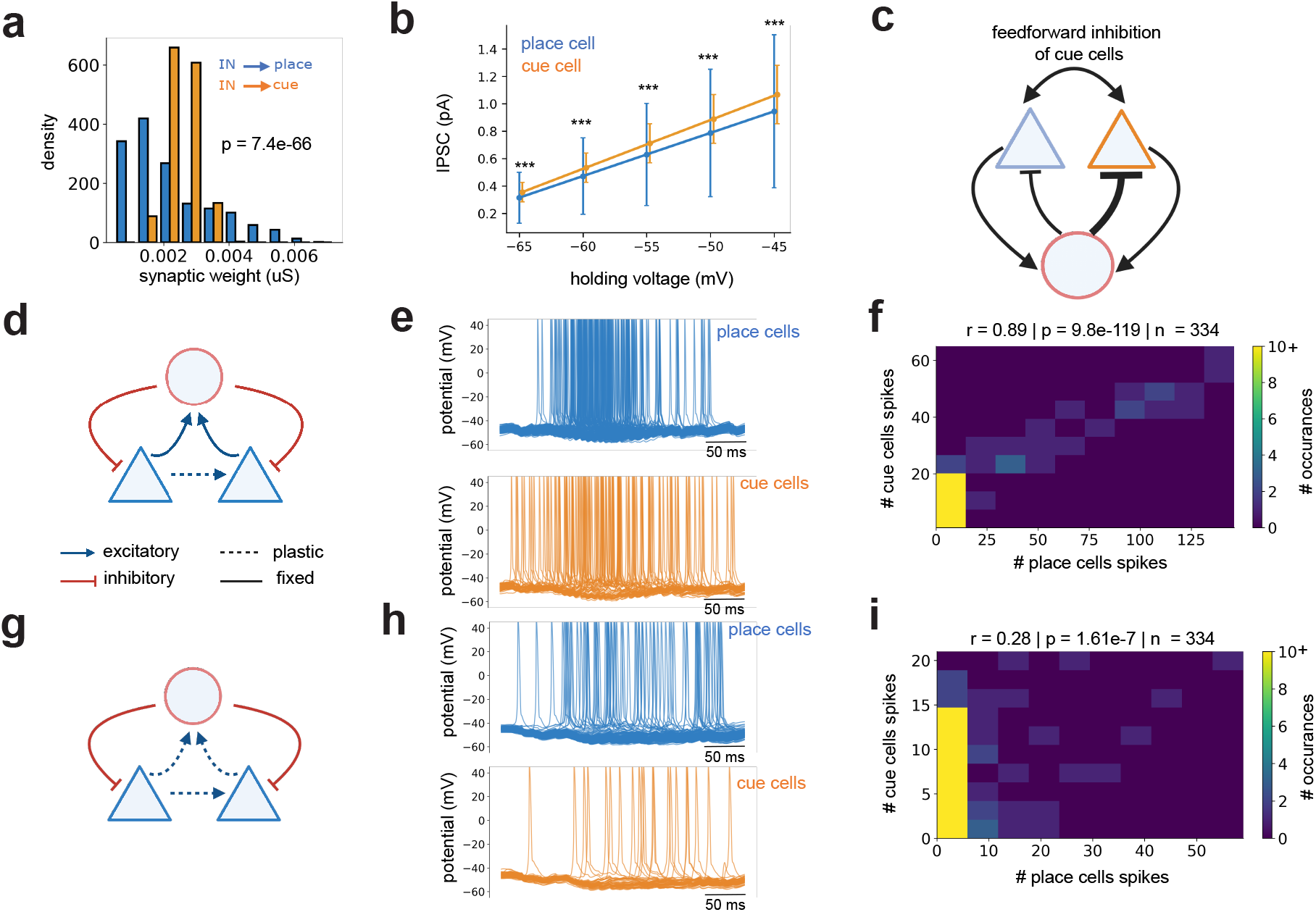
Network perturbations and predictions. **a**. Inhibitory weight onto cue cells is higher than place cells (median *w*_*I*→*cue*_: 0.0025 uS; median *w*_*I*→*place*_: 0.0018 uS; *p* = 7.4 × 10^−66^, one-sided Mann Whitney U-test). **b**. Simulated inhibitory post-synaptic currents (IPSCs) recorded from place cells and cue cells in voltage clamp mode after a single interneuron spiked. Across a range of holding voltages, IPSCs were significantly higher in cue cells (*n* = 960 pairs) than place cells (*n* = 4240 pairs) (one-sided Mann Whitney U-test, \*\*\**p <* 0.001). **c**. Feedforward inhibitory motif: cue cells are more suppressed by interneurons than place cells. **d**. Ablation of inhibitory plasticity: only *E → E* synapses are modified **e**. Somatic membrane potential of place cells (blue) and cue cells (orange) during a spontaneous memory replay event **f**. Heatmap of place-cue cell coupling: when inhibitory plasticity is ablated, the cue cell and the place cell subnetworks are tightly coupled during the offline epoch (Pearson’s *r* = 0.89, *p* = 9.8 × 10^−119^), in contrast to place-cue cell decoupling when inhibitory plasticity was active (Figs 1g, 2g). **g**. Schematic showing the sSTDP rule applied to *E → E* and *E → I* connections. **h**. Somatic membrane potential of place cells (blue) and cue cells (orange) during a spontaneous memory replay event showing that cue cell suppression is lost. **i**. Heatmap of place-cue cell coupling: when *E → E* and *E →I* plasticity is active, the cue cell and place cell subnetworks remain tightly coupled during the offline epoch (Pearson’s *r* = 0.28, *p* = 1.61 × 10^−7^). Hence, *E → I* plasticity is insufficient for cue cell suppression.

Given the observation of powerful feedforward inhibition onto cue cells, we next predicted that targeted ablation of inhibitory plasticity will impair the network mechanism of adaptive stimulus (Terada et al. (2022)). In agreement with the prediction and further supporting the overarching hypothesis that inhibitory plasticity is necessary for world structure learning, models trained with only recurrent *E→E* plasticity exhibited significant co-activity across the cue cell and place cell populations during the offline epoch (Figs 3d-f; Pearson’s *r* = 0.89, *p* = 9.8 *×* 10^−119^). As a control, networks trained with *E→E* plasticity only but containing few cue cells (< 3% of total PC population) were still able to generate spontaneous replay events in the forward and reverse directions (Fig S3 and Fig S4 (top row)) similar to Ecker et al. (2022). Additional simulations reported that restoring inhibitory plasticity restores the adaptive stimulus mechanism (Fig S4 (bottom row)).

To continue testing alternative mechanisms of cue cell suppression, we performed a final simulation in models with *E→E* and *E→I* plasticity (Figs 3g-i). Similar to the model with only *E→E* plasticity (Figs 3d-f), cue cells and place cells were significantly co-active during SPW-Rs (*r* = 0.28, *p* = 1.61 *×* 10^−7^). The collective results of Figs 3d-i and Fig S4 support the hypothesis that inhibitory plasticity is necessary for cue cell suppression. Additionally, from this simulation experiment (Figs 3g-i), we predict that IN firing rates should stay stable during learning, which can be experimentally verified through a multitude of electrophysiological and/or optical methods.

To summarize, we have shown that inhibitory plasticity is both sufficient (Fig 2) and necessary (Figs 3d-i) for cue cell suppression during SPW-Rs. Additionally, we have provided numerous predictions that can be tested experimentally to gather evidence that such a mechanism exists and plays a critical role in memory consolidation. At the cellular level, inhibitory plasticity from *I→E* synapses should lead a powerful feedforward loop onto cue cells that can be measured by increased IPSCs onto cue cells compared place cells (Fig 3a-c). Furthermore, because models trained with *E→E* and *E→I* plasticity did not result in cue cell suppression, IN firing rates should stay stable during learning (Fig 3g-i). At the network level, ablating inhibitory plasticity (Figs 3d-f) should result in loss of cue cell suppression that reemerges by restoring inhibitory plasticity (Fig S4). At the behavioral level, this suggests that ablating inhibitory plasticity would hinder an animal’s ability to perform world structure inference, leading to impairment for goal-oriented behavioral tasks. Taken together, these simulations highlight how synaptic-level mechanisms can cascade to affect network function across multiple spatial scales.

### Simplified model and theoretical analysis

Finally, we provide some intuition about how our model generates spontaneous replay and the role inhibitory plasticity plays in structural learning in the presence of noise.

Consider a maximally abstracted world consisting of a set of “schemas” (which may correspond to physical space, conjunctive context, sensory cues, or abstract concepts). For simplicity, suppose there is a one-to-one correspondence between schemas in the outside world and tuned “concept neurons” in the brain, which we represent with colored circles in Fig 4a. Also suppose a “synapse” exists between any pair of distinct neurons (Fig 4a, weight matrix). The “world structure” to be learned is simply some sequence of schemas.

**Figure 4.**
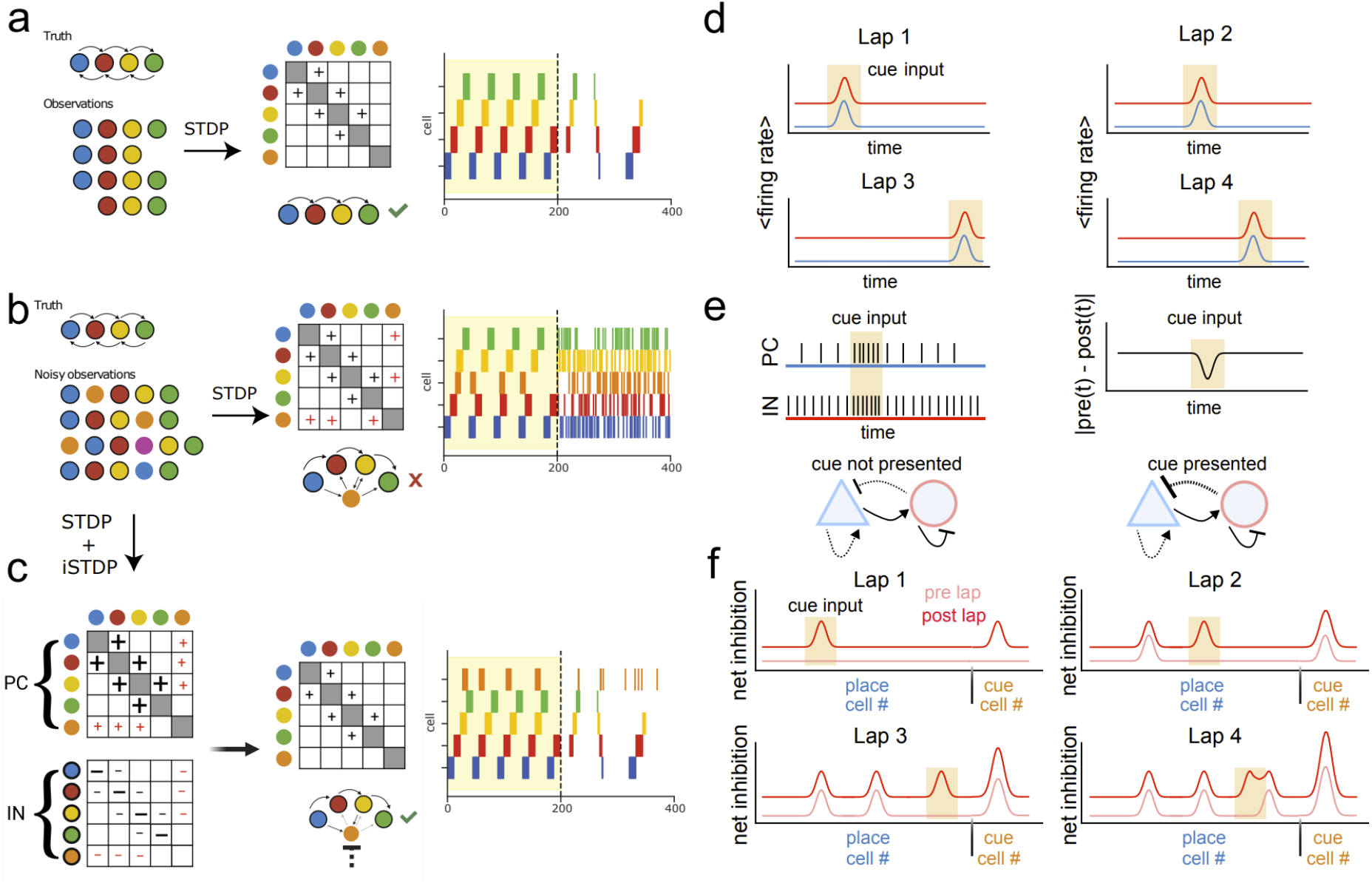
Abstract model of world structure learning in the presence of noise. **a**. World structure (“Truth”) consists of an arbitrary sequence of stimuli to be learned. The “network” consists of “cells” each tuned to exactly one stimulus. In the ideal scenario, observations only consist of subsequences of the ground truth sequence. Symmetric STDP thus bidirectionally potentiates weights along the diagonal of the sorted weight matrix between cells representing adjacent elements of the sequence (right; cf. Fig 1e, left). **b**. A more realistic observation model contains noise: distractor stimuli may appear stochastically at different locations, or sequence stimuli may be erased or appear out of order. As a result, sSTDP potentiates off-diagonal weights (right; cf. Fig 1e, right). Under the realistic observation model of **b**., many off-diagonal synapses are potentiated. Thus, stimulating the first cell in the sequence offline will set off a stochastic cascade, possibly activating side paths through the network via distractor cells. Because distractors appear stochastically, they have positive weight on and may reactivate many other cells, poisoning structured replay. **c**. Active inhibition of these distractor cells rescues replay. As long as distractors are prevented from exciting other cells, the side paths are interdicted while the true sequence undergoes unencumbered reactivation, restoring structured replay. **d**. During each lap along the linear track, a distractor input is driven at a random location (tan boxes). This excitatory input directly target PCs and INs within the local CA3 circuit, therefore increasing the firing rate for both cell populations for the duration that the distractor input is active. **e**. As a consequence of the distractor input increasing firing rates of PCs and INs, it is more likely that the delay between spikes from the PCs and the INs will decrease (i.e., reduction in | *pre*(*t*) − *post*(*t*) |). During this time, the sSTDP rule will generate a larger increase in inhibitory weight from INs onto PCs compared to epochs when the distractor input is turned off. **f**. For each lap, there will be a baseline increase in net inhibition onto place cells and cue cells (*I*_*baseline*_). Additionally, cue cells and place cells that were active during the distractor input will receive enhanced inhibition (*I*_*enhanced*_) for a net inhibitory change of *I*_*baseline*_ + *I*_*enhanced*_. As the same population of cue cells are active at the time the distractor input is presented on each lap, the cue cell population will ’accumulate’ this baseline and enhanced inhibition over *N* laps (*I*_*net,cue*_ = *NI*_*baseline*_ + *NI*_*enhanced*_). In contrast, because the distractor input occurs at a random location on each lap, the enhanced inhibition will be distributed over the place cells, such that each place cell subpopulation will receive enhanced inhibition once over the course of *N* laps (*I*_*net,place*_ = *NI*_*baseline*_ + *I*_*enhanced*_).

To learn this world structure, the network modifies the synaptic weight matrix *W* in response to observations drawn from a generative process parameterized by the ground-truth sequence. To be capable of “world structure learning”, the network must be able to learn any sequence of schemas it is presented with.

In an ideal world, observations only consist of repeated sequential presentations of the ground truth sequence. In this case, a Hebbian LTP rule on excitatory synapses only is sufficient to learn this sequence: every time a transition *a→b* is observed, weights *W*_*ab*_ and *W*_*ba*_ are both incremented by 1. This process potentiates weights along the diagonal between cells representing adjacent elements of the sequence (Fig 4a right; cf. Fig 1e, left). In the offline state, the network receives random input, which may by chance excite an arbitrary neuron *x*_1_ in the network. Without loss of generality, we let *x*_1_ be a one-hot vector (a vector of length *N* with 1 in the position corresponding to the active neuron, zero everywhere else) at the first schema in the sequence. We then run the network for *K* timesteps (where *K* is the number of schemas in the sequence), letting the network follow its internal dynamics *x*_*t*_ = *σ*(*Wx*_*t*−1_), where *σ* is a stochastic nonlinearity that returns a one-hot vector whose nonzero element represents the next active neuron, with probabilities proportional to the *x*_*t*−1_-th column of *W* (see **Methods**). Adaptation in the models of Figs 1, 2 gives sequences “momentum” by preventing network reactivations from sticking in place or doubling back; to model adaptation abstractly, for each event we use only either the lower half-diagonal of *W*, corresponding to the forward weight matrix, *W*_*FWD*_ or the upper diagonal (corresponding to the reverse weight matrix *W*_*REV*_). The output sequence is an offline statistical estimator of the ground truth sequence, which we term REPLAY.

A more realistic observation model contains noise: distractor observations may appear stochastically at different locations in the sequence. Using the previous Hebbian LTP rule, when distractor schemas are observed, off-diagonal weights are potentiated in the weight matrix (Fig 4b right; cf. Fig 1e, right); these off diagonal weights create artificial “side paths” through the network that do not follow the world structure path. Because the stochastic distractor can appear before or after any other schema, it eventually creates potential paths between any two observed schemas in the network as well as cycles at replay time. Since REPLAY samples the weight matrix, aberrant paths in the network translate into replay of disorganized, out-of-order sequences (see also: **Math Supplement**).

To rescue replay in this network, it is sufficient to simply inhibit the distractor neuron (orange) at replay time (Fig 4c (top)). If distractor neurons are not activated, side paths through the network are blocked, and the network dynamics would simply return a walk through the structure of the world. How does the network select only those neurons corresponding to distractors for inhibition compared to neurons encoding part of the structure of the world? We predict that in the online period, the network must represent all schemas because it cannot know *a priori* which schemas will be part of the structure of the world and which will turn out to be distractors. Instead, it must learn selective suppression of schemas that turn out to be distractors, motivating the need for inhibitory plasticity.

To understand why inhibition accumulates on distractor neurons, consider the population firing rate as the mouse navigates the linear track. Because place cells tile the track, the average population firing rate will be near-uniform in the absence of any distractor input. When a distractor input is received by the network, the additional excitatory currents will directly drive both PC and IN populations (see Tables 1-3 for connectivity profiles), causing a local increase in the population firing rates for both cell types relative to the baseline condition of only structured input. As the distractor input moves from lap to lap, the network sees this local increase in firing rate amongst both PC and IN populations at different regions of the track (Fig 4d). This local increase compresses the spike timings of PCs and INs during distractor input, increasing the rate in change of inhibitory weight onto PCs (Fig 4e). This phenomenon potentiates the weights of inhibitory afferents to both place and distractor cells; however, because at each lap a different subset of place cells will be active during the distractor input, the inhibition onto the place cell population is distributed. In contrast, total inhibitory weight onto cue cells is increased in every trial as the same population of cue cells will be activated (Fig 4f).

**Table 1.** Biophysical parameters of local circuit.

|  | PC→PC | PC→INT | INT→PC | INT→INT |
| --- | --- | --- | --- | --- |
| Convergence | 20 | 140 | 20 | 4 |
| Target Compartment | SR/SO-distal | SO | Soma | Soma |
| AMPA rise (ms) | 0.5/0.5 | 0.07 |  |  |
| AMPA fall (ms) | 9.77/8.77 | 0.20 |  |  |
| AMPA conductance (uS) | $3.5e^{-3}$ | $8.0e^{-2}$ | | |
| AMPA reversal (mV) | 0.0 | 0.0 |  |  |
| NMDA rise (ms) | 2.3 |  |  |  |
| NMDA fall (ms) | 100.0 |  |  |  |
| GABA-A rise (ms) |  |  | 0.30 | 0.08 |
| GABA-A fall (ms) |  |  | 6.2 | 4.8 |
| GABA-A conductance (uS) | | | $1.0e^{-3}$ | $1.2e^{-4}$ |
| GABA-A reversal (mV) |  |  | -75.0 | -75.0 |

**Table 2.** Biophysical parameters of afferent connections onto PCs. MF = mossy fiber; MEC = medial entorhinal cortex; LEC = lateral entorhinal cortex.

|  | MF→PC | MEC→PC | LEC→PC |
| --- | --- | --- | --- |
| Convergence | 32 | 20 | 16 |
| Target Compartment | SL/SO-proximal | SLM-proximal | SLM-distal |
| AMPA rise (ms) | 0.5/0.5 | 0.5 | 0.5 |
| AMPA fall (ms) | 144.03/144.03 | 12.66 | 12.66 |
| AMPA conductance (uS) | $7.5e^{-4}$ | $3.35e^{-4}$ | $5.0e^{-3}$ |
| AMPA reversal (mV) | 0.0 | 0.0 | 0.0 |
| NMDA rise (ms) | 2.3 | 2.3 | 2.3 |
| NMDA fall (ms) | 100.0 | 100.0 | 100.0 |

**Table 3.** Biophysical parameters of afferent connections onto INTs. MF = mossy fiber; MEC = medial entorhinal cortex; LEC = lateral entorhinal cortex.

|  | MF→INT | MEC→INT | LEC→INT |
| --- | --- | --- | --- |
| Convergence | 824 | 8 | 8 |
| Target Compartment | SR | SLM-proximal | SLM-distal |
| AMPA rise (ms) | 2.0 | 2.0 | 2.0 |
| AMPA fall (ms) | 6.3 | 6.3 | 6.3 |
| AMPA conductance (uS) | $3.4e^{-4}$ | $1.0e^{-4}$ | $1.35e^{-3}$ |
| AMPA reversal (mV) | 0.0 | 0.0 | 0.0 |

Simulations from our detailed biophysical model support this accumulation of inhibition onto cue cells: we find that inhibitory weights and IPSCs onto cue cells are significantly higher than onto place cells (Figs 3a-c). This observation suggests that the circuit self-organizes to generate powerful feedforward inhibition onto cue cells, which is sufficient to suppress the effect of distractor inputs during electrophysiological events such as ripples.

## Discussion

The precise content of replay, and its role in learning, remains one of the main open questions in the study of the hippocampus. Replay has thus far primarily been studied in highly regular and deterministic experimental environments. However, repeated exposures to the same environment in the real world are not the same: while natural environments contain statistical regularities, these regularities are obscured in imperfect or incomplete observations. A fundamental challenge of learning in the real world is abstracting useful, low-dimensional representations for long-term storage amid a continuous barrage of noisy sensory input (McClelland et al. 1995). Here, we show that a spike-time dependent LTP rule at CA3 GABAergic synapses is sufficient to enable consolidation of the statistical structure of the world from noisy experience. Using a three-pronged approach of spiking network, biophysical, and normative modeling, we demonstrate that this mechanism is biologically plausible, interpretable, and reproduces experimental findings. We make the major prediction that the ablation of plasticity at GABAergic synapses will impair an animal’s ability to filter irrelevant stimuli out of replay. Finally, we simplify the intuitive notion of “statistical learning” by the hippocampus into a statistical inference problem that must be solved using biological primitives, and argue mathematically that the symmetric STDP-based biological algorithm we propose correctly solves this inference problem in the asymptotic limit of observations.

The prevailing hypotheses about replay content hold that replay reinstates either predictive representations or specific past experiences. While replay content reflects planning in some tasks (Ólafsdóttir et al. 2018, Foster 2017), it is divorced from imminent plans in others (Gillespie et al. 2021, Singer & Frank 2009, Ambrose et al. 2016, Karlsson & Frank 2008). These non-immediately useful replay events may nonetheless be predictive if replay stores and organizes specific experience for later use (Gillespie et al. 2021, Wittkuhn et al. 2021). The specific-experience model predicts that replay events should faithfully reactivate sequences representative of individual experiences of the environment. However, the findings of Terada et al. (2022), through the lens of the results presented here, challenge this model as well: even when prominent random stimuli dominate the animal’s specific experiences, representations of stimuli that correspond to unique experiences are not only not reactivated but suppressed from replay events.

Here, we propose a model-based replay framework which we believe shows that these viewpoints are not mutually exclusive. We hypothesize that replay neither reprises specific past experiences, nor generates concrete predictions or plans of future behavior, but rather returns samples from the animal’s working model of the deeper structure of its world. This conceptual model synthesizes elements of both of these theories to explain how replay can be useful for planning, even though representations from distal past experience, which are not imminently behaviorally relevant, are sometimes replayed. The ability to draw samples from a distribution is computationally powerful, as it enables downstream circuitry to compute arbitrary queries on the underlying model. For instance, these samples are sufficient for a downstream brain area to implement the simplest model-based reinforcement learning strategy, random shooting(Richards 2005), with minimal complexity. We note also that this framework is agnostic to the *semantics* of the replayed representations, and thus has explanatory and predictive power irrespective of whether the underlying experience consists of spatial, auditory, social, or more general abstract sequences and relationships. Our plasticity rule simply embeds the topology of a world into the weight matrix of a randomly recurrently connected network. The networks in our simulations were presented with 1D sequences, mirroring the sequential nature of experience in any environment: for a 2D or more complex environment, the world must still be experienced incrementally, through sequential trajectories through it. Recent work has shed light on the spontaneous organization of place “schemas” in the hippocampus (Bittner et al. 2017, Rolotti et al. 2022, Priestley et al. 2021); once such representations are organized, the *A→B* relationships between them can be learned incrementally even if gross sequences are not repeated. Such a learning rule allows previously unobserved sequences to be hypothesized and replayed in the brain. We predict that world structures which are fundamentally *non*-spatial, corresponding to arbitrary relational rules, can also be learned and replayed by the hippocampus via this mechanism.

Much work has been done to characterize Hebbian-type learning rules in the cortex. This work has primarily focused on glutamatergic synapses onto principal cells (Hebb 1949, Markram et al. 1997), and to a lesser extent glutamatergic synapses onto interneurons (Kullmann & Lamsa 2007). The traditional conceptual model does not include plasticity at the GABAergic synapse, under the presumption that inhibition provides a constant background for learning. Nonetheless, long-term plasticity of GABAergic synapses has previous been reported in hippocampal slices(Gaiarsa et al. 2002), although whether the predominant effect is potentiation(Caillard et al. 1999*a*, Nusser et al. 1998), depression(Chevaleyre & Castillo 2003, Wang & Stelzer 1996, Stelzer et al. 1987, 1994, Caillard et al. 1999*b*, 2000, Loisy et al. 2022), or some combination(Gubellini et al. 2001) requires clarification *in vivo*. Compellingly for our work here, D’amour & Froemke (2015) showed a symmetric spike time-dependent LTP rule for the potentiation of inhibitory synapses in auditory cortex, though the computational function of this rule remains unclear. Our results indicate that the phenomenon of world-structure replay is robust for a family of inhibitory LTP rules (Figs S1, S2), providing further support for the biological plausibility of this rule.

A limitation of this work is that we have treated interneurons as a uniform population. In reality, distinct subtypes of interneurons play very different roles computational roles in the hippocampus (Royer et al. 2012, Pelkey et al. 2017, Geiller et al. 2020, Klausberger & Somogyi 2008). The size of the interneuron population required to achieve this effect in a spiking network of thousands of neurons is comparatively small—therefore, it is possible that the inhibitory plasticity mechanism we describe is localized to a particular subtype. Our model predicts that the relevant interneuron population will respond to cues during exploration and will be recruited to SPW-Rs at rest. Dissecting the subtypes shaped by this inhibitory LTP rule represents an important avenue for future experimental work.

In many hippocampal-dependent tasks, certain stimuli carry intrinsic valence, such as the delivery of a reward, which may shape replay statistics via neuromodulation in ways we have not considered here. Previous work has suggested alternative interpretations of the hippocampal place code through this lens, e.g. as a predictive map (Stachenfeld et al. 2017); we speculate that an inhibitory plasticity mechanism may also be useful for learning such representations in the presence of noise. Our inhibitory plasticity mechanism supports both forward and backward denoised replay, though other work has suggested differing functions for different directions (Mattar & Daw 2018), which could be incorporated as an instructive signal into our model based on cognitive demands.

We have provided a mechanistic narrative of how the hippocampus may incrementally assemble a “clean” internal representation of the hidden structure of the world despite only ever receiving noisy observations. The concurrent demands of expressiveness and flexibility provide insight into why a plasticity mechanism, in particular one of inhibitory synapses, is necessary for robust learning in the face of distractors—to wit, the brain cannot know *a priori* which stimuli are important until it has sufficiently sampled the environment. The same stimulus which is a distractor in one context may be essential to understanding the structure of the world in the next. Therefore, the hippocampus must be able to robustly represent all stimuli during online learning as they are received, then selectively consolidate only those representations which model the conserved underlying structure of the ensemble of its experience. We make a number of novel, experimentally-approachable predictions using this framework that will hopefully provide insight into the low-level mechanisms underlying high-level cognitive functions such as world structure inference. We believe that the framework we have presented here provides a bridge between the synaptic and the cognitive levels of our understanding of learning, and if experimentally validated, could shed light on the fundamental question of how distributed, local learning rules can structure initially random neuronal networks to perform highly complex cognitive tasks.

## Acknowledgments

Z.L. is supported by NINDS 1F31NS120783-01 and NIGMS 5T32GM007367-45. D.H. is supported by American Epilepsy Society Predoctoral Research Fellowship and NINDS 1U19NS104590. I.S. is supported by NINDS 1U19NS104590. A.L. is supported by NIMH 1R01MH124047, 1R01MH124867; NINDS 1U19NS104590, 1R01NS121106, and the Kavli Foundation. We thank Drs. L. Abbott, S. Fusi, and C. Schevon for their comments on previous versions of the manuscript.

## Author Contributions

All authors conceptualized the study. Z.L. and D.H. performed computational simulations, mathematical analysis, and data visualization. All authors wrote the manuscript.

## Lead Contact and Materials Availability

Further information and requests for resources and reagents should be directed to the Lead Contact Attila Losonczy. All unique resources generated in this study are available from the Lead Contact with a completed Materials Transfer Agreement. Code is available at.

## Method Details

### Spiking network model

We first built a random network model of a CA3 subnetwork capable of generating spontaneous replay, similar to previous work (Ecker et al. 2022) (see reference for detailed equations and constants). All network simulations were run in Brian2 (Stimberg et al. 2019).

### Neurons

We initialized the network with 8000 pyramidal cells and 150 interneurons. Each neuron was modeled by a variant leaky integrate-and-fire model with cellular adaptation (Ecker et al. 2022, Gerstner et al. 2014).

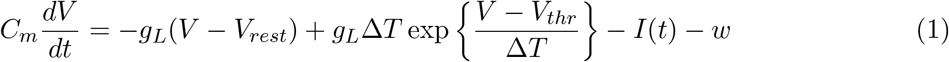

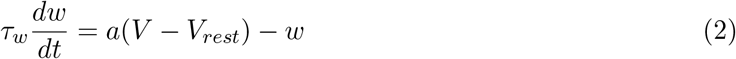

where *C*_*m*_ is membrane capacitance, *g*_*L*_ is the leak conductance, *V*_*rest*_ is the resting potential, *V*_*thr*_ is the spiking threshold, Δ*T* is a shape parameter, *w* is the adaptation current and *I* is the input (synaptic) current. When *V* reaches the threshold *V*_*thr*_, a spike is recorded and *V* is reset to a hyperpolarized reset potential *V*_*reset*_, where it is held for a refractory period *t*_*ref*_. Adaptation strength is given by *a*.

### Synapses

Synapses were modeled as double-exponential time-varying conductances

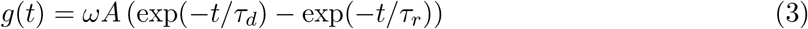

where *ω* is the weight of the synapse, *τ*_*r*_, *τ*_*d*_ are rise and decay time constants. The postsynaptic current was modeled with excitatory and inhibitory components as in (Ecker et al. 2022) as

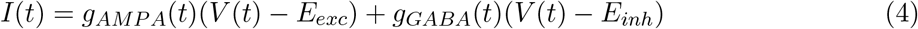

where *E*_*exc*_ = 0, *E*_*inh*_ = −70 mV.

### Network topology

Let *ℐ* be the set of inhibitory neurons and *ℰ* be the set of pyramidal neurons in the network. A random network topology was selected by sampling a directed Erdős-Rényi graph *G*(8000, 0.1) (Erdős & Rényi 1959): i.e., a synapse between each (ordered) pair of pyramidal neurons (*i, j*) *∈ ℰ × ℰ* exists with probability 0.1 (drawn i.i.d.). The *I→E* topology was determined using a similar process: each ordered pair (*i, j*) *∈ ℐ × ℰ* exists as a synapse in the network with probability 0.25.

### Training phase

In the online phase, the network receives a series of observations and modifies its synaptic weights accordingly in order to estimate the “world structure” sequence that gave rise to those observations. The simulated behavior trace was generated as a simulated animal moving at constant velocity through a circular 1D track.

Simulated spike trains were sampled from a Poisson process with a baseline firing rate of *λ* = 0.1 Hz.

A random 50% of pyramidal cells were selected as place cells and received spatially tuned input in a preallocated place field covering 10% of the track, tuned such that the mean firing rate in the center of its place field was 20 Hz. The firing rate *λ*_*i*_(*x*) of cell *i* at each spatial position *x* was given

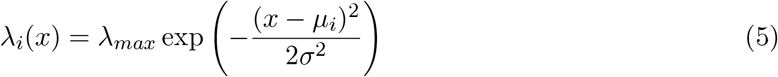

where *µ*_*i*_ is the peak of cell *i*’s tuning curve and *σ* parameterized.

A random 15% of neurons were chosen to be “cue cells”, consistent with findings by (Terada et al. 2022). A cue input was presented at a random location at each lap. The firing rate *λ*(*x*) of all cue cells at each spatial position *x* on lap *k*

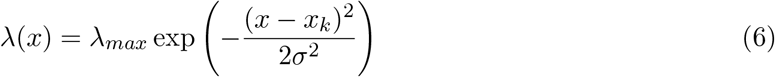

where *x*_*k*_ corresponds to the location of cue presentation on lap *k*

Spikes within a 5 ms refractory period of a previous spike were rejected.

A symmetric STDP rule was used to train the network as follows: For each synapse *i→j* with weight *w*, for each pair of spikes

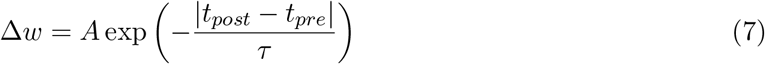

This rule was used at both excitatory and inhibitory synapses, with different values of *τ*_*E*_, *τ*_*I*_.

To explore the space of inhibitory plasticity rules, we examined the family of spike-time dependent plasticity rules of the form (Gerstner et al. 2014)

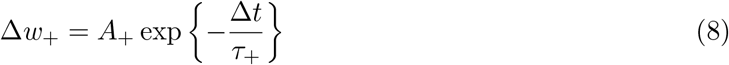

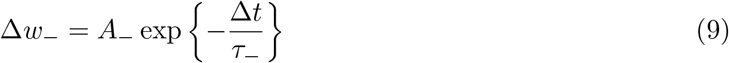

where Δ*w*_+_ is applied to the synapse if *t*_*pre*_ *< t*_*post*_ (causal spiking), Δ*w*_−_ is applied to the synapse if *t*_*pre*_ *> t*_*post*_ (anticausal spiking). We varied the varying the parameters *A*_+_, *A*_−_, *τ*_+_, *τ*_−_. When *A*_+_ = *A*_−_ *>* 0, we obtain the standard symmetric STDP rule. When *A*_−_ = −*A*_+_, we obtain the classic (asymmetric) STDP rule. Varying *τ*_+_, *τ*_−_ independently while holding *A*_+_ = *A*_−_ *>* 0 gave a variety of LTP rules. Throughout this exploration, we held the *E→E* plasticity rule fixed as the experimentally demonstrated rule (Mishra et al. 2016)

### Offline (replay) phase

In the offline state, excitatory cells received random low-level (0.1 Hz) Poisson spiking input. Replay emerged spontaneously in the network.

### Biophysical model

The biophysical model of hippocampal CA3 was built from 260 CA3 pyramidal cells (PCs) and 30 fast-spiking perisomatic-targeting interneurons (INs). Each neuron was represented as a multi-compartment model, where each compartment mapped to various physical locations including soma, basal dendrites, and apical dendrites. Additionally, each neuron contained biophysical ion channel models with realistic conductances, thereby allowing the user to track the continuous intracellular voltage trace within each compartment. PCs and INs received external input from multiple afferent sources modeled as Poisson spike generators, including the medial entorhinal cortex (MEC), lateral entorhinal cortex (LEC), and the dentate gyrus (DG) via mossy fibers (MFs).

The neurons were placed into a virtual CA3 volume divided into stratum lacunosum moleculare (SLM), statrum radiatum (SR), stratum lucidum (SL), stratum pyramidale (SP) and stratum oriens (SO). The soma position of PCs and INs as well as specific connectivity patterns between PCs, INs, and external inputs (convergence and specific site of contact) were allocated according to know experimental data. For further details on biophysical parameters, see Tables 1-3.

### Training and replay

Similar to the spiking network model, a virtual circular track was generated and simulations were performed with the mouse moving at a constant velocity. All simulations were run in the NEURON environment (Hines & Carnevale 2001) with *dt* = 0.1 ms.

For training, 80% of CA3 pyramidal cells were selected as place cells and received preallocated and spatially tuned input from external MEC and MF sources. Specifically, MFs carried place inputs while MEC afferents carried grid inputs. The diameter of place fields was selected to cover 10% of the track, tuned such that the mean firing rate at the center of the place field was 20 Hz (see spiking network model for equations). Grid inputs were instantiated with similar parameters but were additionally allowed to cover the virtual track in a periodic manner with a grid spacing equal to 10% of the track. The remaining 20% of CA3 pyramidal cells were selected as cue cells and received preallocated distractor input from external LEC. The distractor input was activated at a random location once per lap.

During the offline epoch, we removed spatially structured and distractor input from all external sources. Instead, CA3 neurons received random Poisson spiking input, and spontaneous replay was observed.

### Quantitative analysis

#### Offline cue-place aggregate analysis

The full offline epoch was divided into non-overlapping bins equating to 100 ms of simulation time. The number of spikes for cue cells and place cells in any given bin was extracted and plotted on a 2D histogram. Pearson’s r was quantified and a two-tailed t-test was used to determine if the cue and place cell sub-networks were functionally connected (p < 0.05) or not (p > 0.05).

#### Abstract model

Let *𝒳* be a finite set of schemas. Define a *sequence* of length *K* as a string *X* = (*x*_1_, *x*_2_, *x*_3_, …, *x*_*K*_) with each *x*_*i*_ *∈ 𝒳* and *x*_*i*_ ≠ *x*_*j*_, *i* ≠ *j*. Let “world structure” consist of a ground truth sequence *Y* = (*y*_1_, *y*_2_, *y*_3_, …, *y*_*K*_) *∈ 𝒳* ^*K*^. For simplicity, we consider only forward replay; the analysis for backward replay is identical.

Real observations consist of a mix of fixed elements of the world and experience-specific stimuli. Therefore, we draw *observations X* i.i.d. according to the following generative process: for each (1 *≤ i ≤ K* − 1), let *x*_*i*_ = *y*_*i*_ (ground truth) with probability 1 − *p* and *x*_*i*_ = *z* (distractor) with probability *p*.

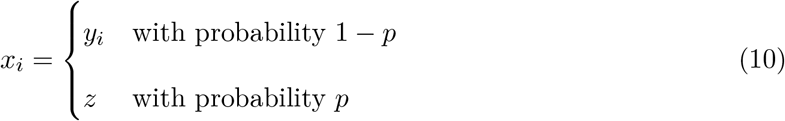

Assume each character *x ∈ 𝒳* is represented by a single “neuron”, and let *W* = |*𝒳* | *×* |*𝒳* | be a matrix initialized with zeros representing synaptic weights between these neurons. Assume for the sake of simplicity that the network is fully connected (equivalently, let each model neuron represent the equivalence class of neurons coding for each stimulus).

Let *W*^*I*^ be another |*𝒳* | *×* |*𝒳* | matrix representing feedforward inhibitory weights from the subset of inhibitory neurons tuned to inputs onto *x*.

Now we introduce the plasticity rule. Under the simplest model of inhibitory plasticity, symmetric LTP of inhibitory synapses occurs via STDP. For each transition *x*_*i*_*→x*_*i*+1_ in an observation, increment 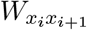 by 1. Increment the inhibitory weight 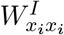 by 1, and the weights 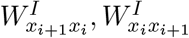 by *β <* 1, which parameterizes the inhibitory plasticity kernel decay rate.

### Replay phase

In the offline phase, the network reads out its running estimate of the “world structure” sequence.

Let *ŷ*_*t*_ be a one-hot vector representing the neuron active at time *t* in a replay sequence. Then *W ŷ* denotes the (unnormalized) distribution of neurons active at time *t* + 1.

We add the inhibitory term −*αW*^*I*^*ŷ*, where *α* is a parameter controlling the relative strength of inhibition and excitation. As inhibition predominates during replay, we pick *α >* 1: for instance *α* = 1.5. Finally, to model stochasticity in sequence activation, we introduce a stochastic nonlinearity: For a nonnegative vector *w*, let the random variable *σ*(*w*) denote the random one-hot vector obtained by drawing a sample from the categorical distribution with probabilities proportional to *w* (if any values of *w* are negative, we replace them with 0).

Hence, we can write the dynamics of the network during replay as:

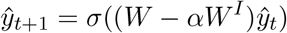

### The REPLAY estimator

Without loss of generality, let *ŷ*_0_ = *y*_0_ (i.e., the first element in the ground-truth sequence). Run this network for *K* timesteps. We define the REPLAY estimator *Ŷ* to be

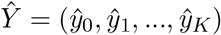

## Data and Software Availability

Software from this study is available at https://losonczylab.github.com.

## Supplementary Information

**Figure S1.**
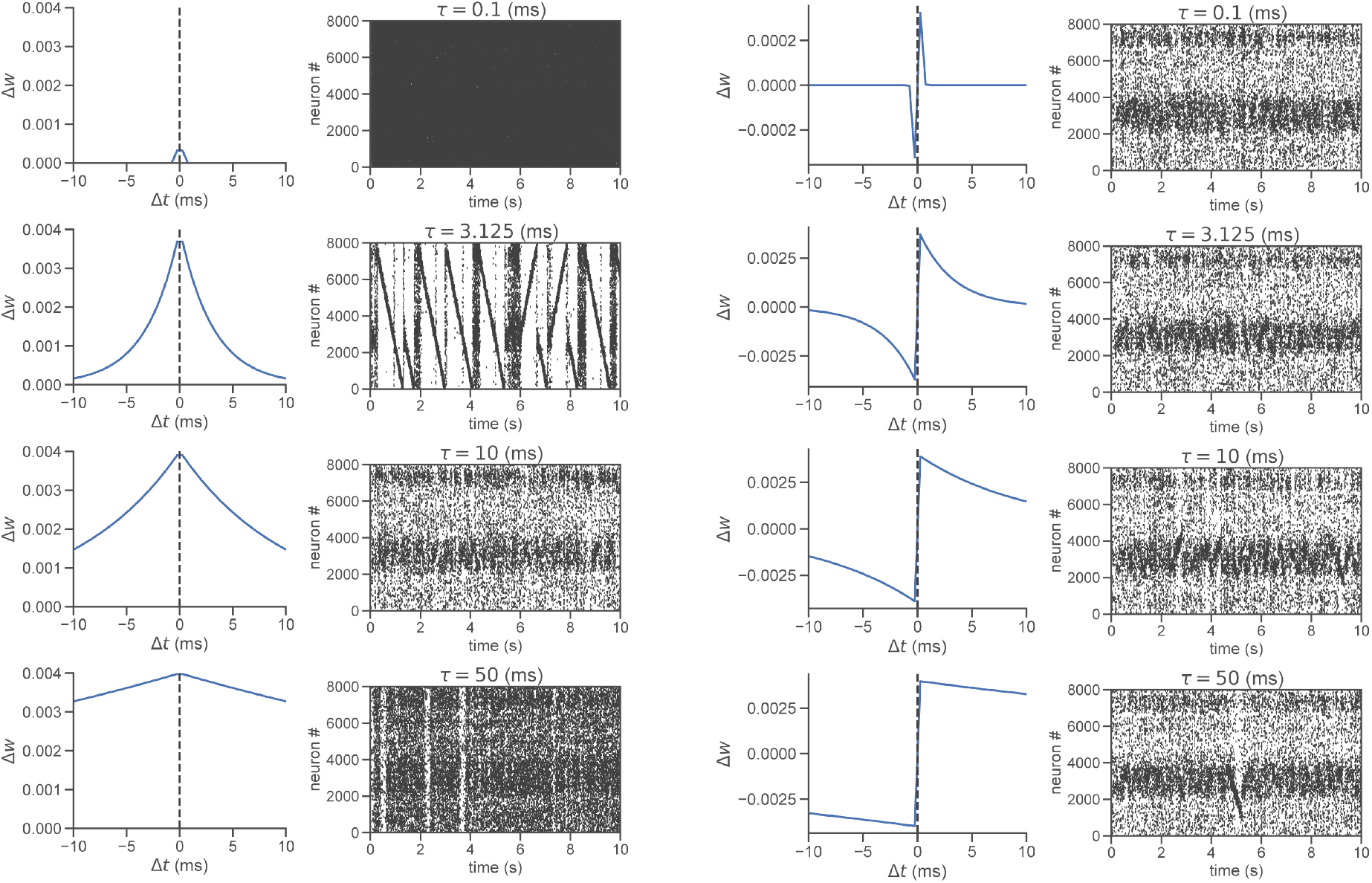
World structure replay under different symmetric and antisymmetric LTP and LTP-LTD rules. Left: Various symmetric STDP kernels. Right: No replay using classic (i.e., antisymmetric) interneuron STDP

**Figure S2.**
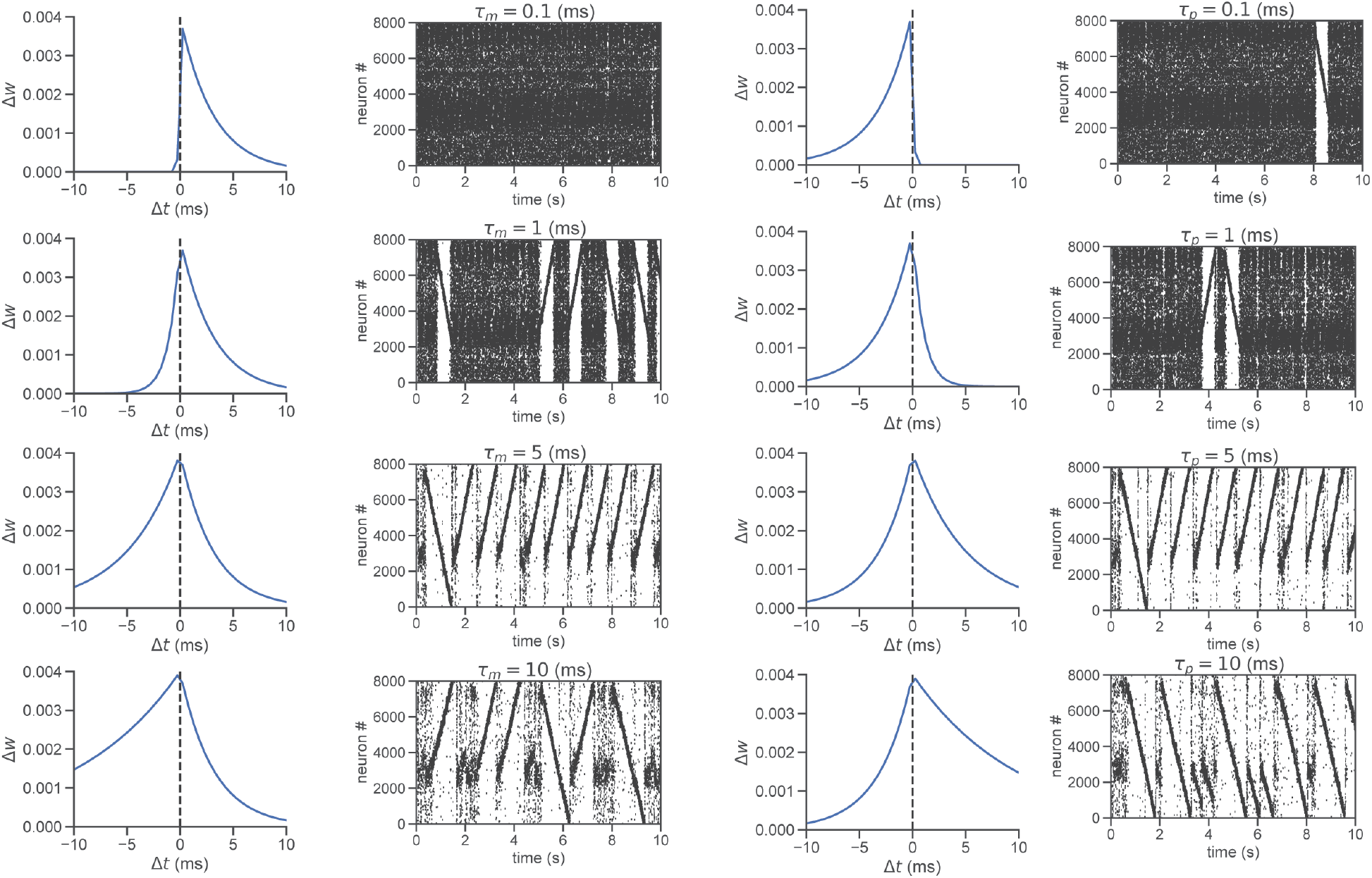
Robustness of world structure replay to different asymmetric Hebbian LTP-type GABAergic plasticity rules. We examined STDP rules defined by a variety of asymmetric LTP kernels and find broadly the same results.

**Figure S3.**
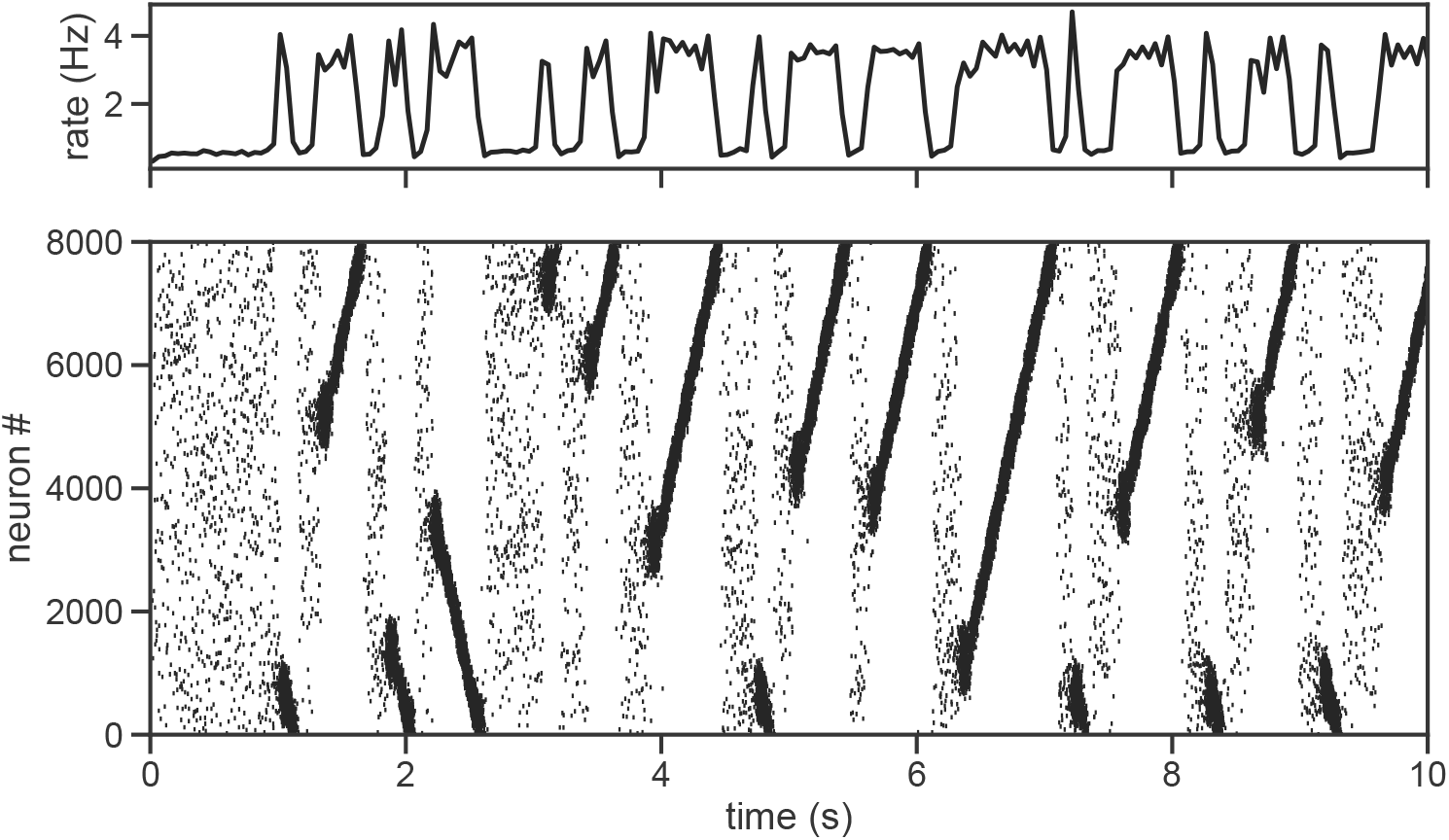
Network without distractor input. In the absence of distractor input, spontaneous replay occurs even without inhibitory plasticity.

**Figure S4.**
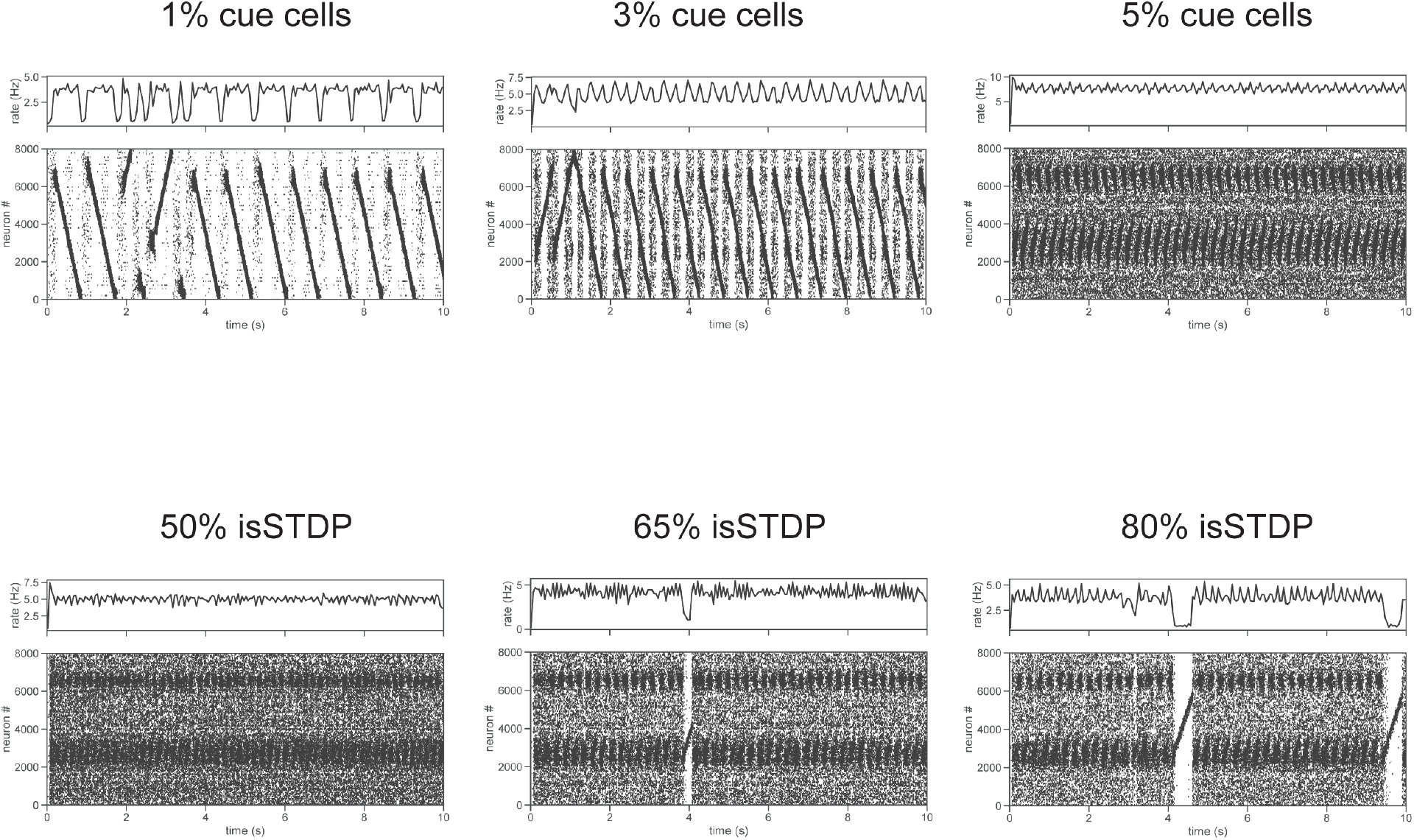
Varying degree of distractor input and inhibitory plasticity. Top: Titrating proportion of distractor (“cue”)-tuned cells in the absence of inhibitory plasticity. Replay is abolished in the network with even 5% distractor cells. Bottom: Titrating inhibitory plasticity weight (relative to that in Fig 1): at 50% or less, replay is abolished altogether.

## Math supplement: The REPLAY estimator

### Assumptions

Let *𝒳* be a finite set. Define a *sequence* of length *K* as a string *X* = (*x*_1_, *x*_2_, *x*_3_, …, *x*_*K*_) with each *x*_*i*_ *∈ 𝒳* and *x*_*i*_ ≠ *x*_*j*_, *i* ≠ *j*.

Let “world structure” consist of a ground truth sequence *Y* = (*y*_1_, *y*_2_, *y*_3_, …, *y*_*K*_), where each *y*_*i*_ *∈ 𝒳*. For simplicity, we will perform this analysis on forward replay only; the analysis for backward replay is identical.

### Online (learning) phase

In the online phase, the network receives a series of observations and modifies its synaptic weights accordingly in order to estimate the “world structure” sequence that gave rise to those observations.

Real observations consist of a mix of fixed elements of the world and experience-specific stimuli. Therefore, draw *observations X* i.i.d. according to the following generative process: for each (1 *≤ i ≤ K* − 1), let *x*_*i*_ = *y*_*i*_ (ground truth) with probability 1 − *p* and *x*_*i*_ = *z* (distractor) with probability *p*.

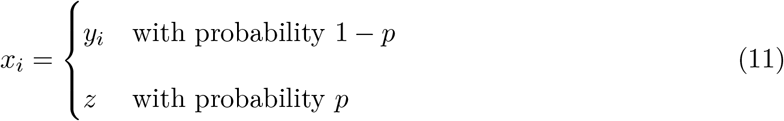

We will perform the analysis assuming one distractor stimulus, though it is straightforward to extend it to multiple distractors.

Now we introduce the plasticity rule. Under the simplest model of inhibitory plasticity, symmetric LTP of inhibitory synapses occurs via STDP. For each transition *x*_*i*_*→x*_*i*+1_ in an observation, increment 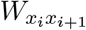 by 1. Increment the inhibitory weight 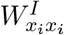 by 1, and the weights 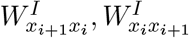 by *β <* 1, which parameterizes the inhibitory plasticity kernel decay rate.

### Offline (replay) phase

In the offline phase, the network reads out its running estimate of the “world structure” sequence.

We add the inhibitory term −*αW*^*I*^*ŷ*, where *α* is a parameter controlling the relative strength of inhibition and excitation. As inhibition predominates during replay, we pick *α >* 1: for instance *α* = 1.5.

Finally, to model stochasticity in sequence activation, we introduce a stochastic nonlinearity: For a nonnegative vector *w*, let the random variable *σ*(*w*) denote the random one-hot vector obtained by drawing a sample from the categorical distribution with probabilities proportional to *w* (if any values of *w* are negative, we replace them with 0).

Hence, we can write the dynamics of the network during replay as:

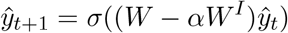

### The REPLAY estimator

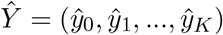

#### Lemma 1.

*Given perfect observations and no inhibitory plasticity, the REPLAY estimator is consistent*.

*Proof*. Each trial consists of an observation of the ground truth sequence *y*_1_*→y*_2_*→ … →y*_*K*_.

Therefore, after *N* trials, the *y*_*i*_, *y*_*j*_ entry of *W* will be *N* if *i* = *j* + 1, and 0 otherwise. Thus, if *ŷ*_*t*_ = *y*_*t*_, then *W ŷ*_*t*_ is simply the *y*_*t*_th column of *W*, which is *N* in the entry corresponding to *y*_*t*+1_ and 0 everywhere else; therefore, for any *N >* 0, *ŷ*_*t*+1_ = *y*_*t*+1_. By induction, we conclude that this estimator is consistent. ▪

#### Lemma 2.

*Given noisy observations (as in Equation 11) and no inhibitory plasticity, the REPLAY estimator is not consistent*.

*Proof*. Consider the *y*_*t*_ column of *W*. Given the Bernoulli generative process from Equation 11, 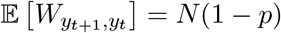 (i.e., the entry corresponding to a transition *y*_*t*_*→y*_*t*+1_ is *N* (1 − *p*) in expectation), while 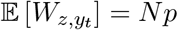. As *N →∞*, the true entries converge to these expected values by the law of large numbers.

Now consider a ground-truth sequence of length *K*: the probability of recapitulating that sequence in this network using the REPLAY estimator is:

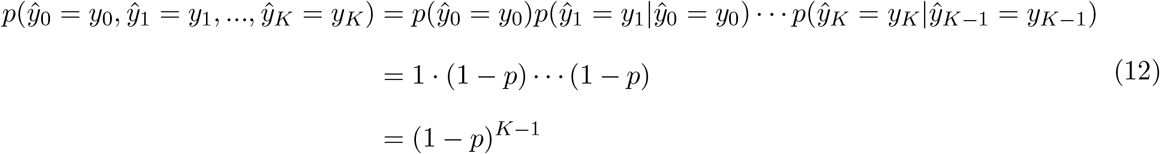

Equivalently, this probability is the probability of the estimator never outputting *z*, where by assumption, *p*(*ŷ*_0_ = *y*_0_) = 1, and in general for *t >* 0, *p*(*ŷ*_*t*_ = *y*_*t*_|*ŷ*_*t*−1_ = *y*_*t*−1_) = 1 − *p*. Thus, as long as *p >* 0, *p*(*Ŷ /*= *Y*) is bounded away from 0 even as *N →∞*. ▪

We make two observations: first, if *p <* 0.5 the ground truth sequence is the single most likely sequence to be replayed. However, absent any other mechanism, because of *K* in the exponent, the probability of replaying this sequence decays quite quickly even for conservative values of *p, K*: for instance, when *p* = 0.1, *K* = 7 we already have *p*(*Ŷ* = *Y*) *<* 0.5 – in reality, the probability of observing a distractor may be significantly higher than 0.1, and much longer sequences may have to be replayed. Therefore, some other mechanism must act to actively suppress distractor replay.

#### Theorem 3.

*For appropriate choice of plasticity kernel width and inhibitory strength, the REPLAY estimator with inhibitory plasticity is consistent even in the presence of noise*.

*Proof*. Just as in Lemma 2, we consider the *y*_*t*_ column of *W*, with 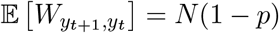 and 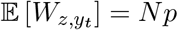. Now we consider the same column of *W*^*I*^: the entry 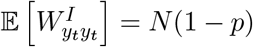, while the entries 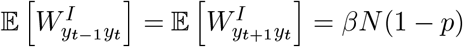. Finally, the entry 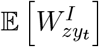 is *β×* the expected number of times *y*_*t*_ is preceded or followed by *z*, i.e. 2*βNp*.

Once again appealing to the law of large numbers, as *N →∞*, the *y*_*t*_th column of *W* − *αW*^*I*^ contains *N* (1 − *p*)(1 − *αβ*) in the position corresponding to *y*_*t*+1_ and *Np*(1 − 2*αβ*) in the position corresponding to *z*, and zeroes elsewhere. Now if 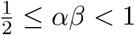, we will have 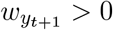 and *w*_*z*_ = 0, which guarantees that *ŷ*_*t*+1_ = *y*_*t*+1_. This proves that the REPLAY estimator with inhibitory plasticity is consistent. ▪

For any *α, β, p* we can calculate the exact probability *p*(*ŷ*_*t*+1_ = *y*_*t*+1_|*ŷ*_*t*_ = *y*_*t*_) in the asymptotic *N* limit as

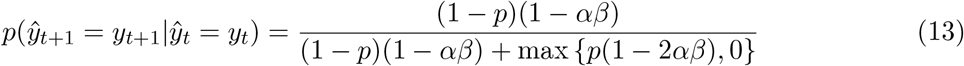

## Math supplement: Realizability in real networks

In real networks, each concept or “schema” is represented by more than one neuron, and the number of neurons is finite. Therefore, it is natural to ask in what parameter regime the world structure sequence is even realizable–for the ground truth sequence *y*_1_*→y*_2_*→ … →y*_*K*_ to be learned and replayed, there must at minimum exist a chain of monosynaptically connected neurons tuned to *y*_1_, *y*_2_, …, *y*_*K*_. This requirement may be relaxed with recurrence (i.e., *y*_1_-tuned neuron *a* excites a second *y*_1_-tuned neuron *b* which in turn excites a *y*_2_-tuned neuron), but we can show that this is not necessary in many cases.

We will now analyze our spiking network model of CA3 from Fig 1. We modeled CA3 as an Erdős-Rényi graph *G*(*N, p*) (Erdős & Rényi 1959): in brief, let *N* be the number of neurons in a network, and 0 *≤ p ≤* 1. *G*(*N, p*) is a generative process defined on graphs *G* = (*V, E*) with |*V* | = *N* nodes such that for any pair of nodes *i, j ∈ V*, the edge (*i, j*) *∈ E* with i.i.d. probability *p*.

Suppose there are *K* schemas represented in this network, where each neuron is tuned to 1 schema, such that schema #1 is represented by *N*_1_ neurons *V*_1_, schema #2 by *N*_2_ neurons *V*_2_, etc., and

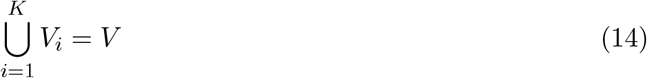

For simplicity, assume *N*_1_ = *N*_2_ = *…* = *N*_*K*_ = *N/K* (where schemas are assigned to neurons uniformly at random), and *V*_*i*_ *∩ V*_*j*_ = *∅* for *i /*= *j*. This is a coloring of *G*(*N, p*) where each “color” corresponds to a schema. Call the parameter *n* = *N/K* the *representational redundancy* of the network, and its reciprocal 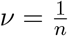. the *representational density*.

We have the following theorem

### Theorem 4.

*Length-K sequences are realizable with high probability in a network of O*(*K*) *neurons*.

*Proof*. Let *V*_*i*_ be the cells tuned to the schema *y*_*i*_ and 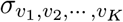 be the indicator random variable

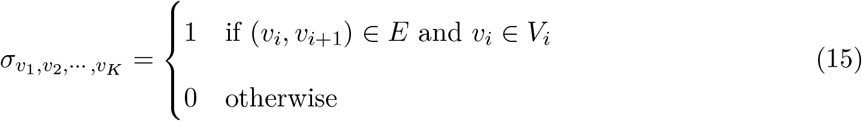

Then the expected number of chains (paths) through the network consistent with sequence *y*_1_*→*…*→y*_*K*_ is

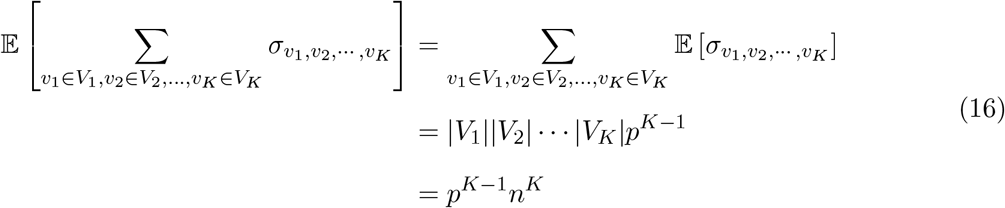

where the second equality follows from the linearity of expectation.

This expectation is greater than 1 when *N > K/p · p*^1*/K*^ = *O*(*K*). ▪

This theorem shows that the number of neurons required to realize sequences of *K* schemas scales only linearly with *K*, rather than quadratically, exponentially, etc. A supralinear scaling rate would impose a sublinear ceiling on the lengths of realizable chains, which may be short in practice given the finite size of the network, thus limiting replay to short sequences. While we do not rule out the existence of such a limit due to other factors, the random topology of the network does not inherently impose such a limit.

### Corollary 5.

*As K becomes large, length K sequences are realizable in the network only if p > ν*.

For the above analysis we treated the connectivity parameter of the network *p* as a constant. However we observe from Equation 16 the expression

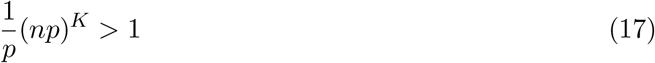

This expression illustrates the realizability of sequences in a random network in terms of the network’s representational redundancy and connectivity—as *K* becomes large, length-*K* sequences are realizable only if *np >* 1 ⇒ *p > ν*; otherwise (*np*)^*K*^ *→*0. In other words, consistent with intuition, networks with more representational redundancy can afford to be more sparsely connected, while networks with sparser *representations* require denser connectivity to achieve the same level of expressiveness in realizable sequences.

## Math supplement: Storing multiple representations

Sometimes, multiple sequences built from the same schemas in a different order must be stored in the same network simultaneously. There are *K*! such sequences possible (assuming every schema is used). A natural question is, how many such sequences can be stored simultaneously while avoiding ambiguity, i.e., one memory sequence corrupting another via use of the same cells. This is equivalent to the question of, how many walks can be drawn in given coloring of *G*(*N, p*) which touch each color exactly once (each in different order) and use each neuron at most once.

We note that this is distinct from the case where such reuse may actually be helpful to learning world structure, e.g. in an open-field maze with multiple approaches to the same spatial location, it may be helpful to reuse that representation across multiple memories of exploration in the maze to enable simulation of trajectories not yet taken.

There are two cases it is relevant to consider: a “best case” where sequence representations are deliberately chosen to avoid reuse of cells, and an “average case” where sequences are chosen randomly. We will consider these cases separately.

### Best case performance

In the proof of Theorem 4, we showed that given a fixed sequence, in expectation there are *p*^*K*−1^*n*^*K*^ chains through the network satisfying that sequence. Suppose we have stored such a sequence in the network already–then there are *n* − 1 neurons in each ensemble left for subsequent chains while avoiding reuse. Let *k* be the maximum number of sequences we can stored in the network. Then we have

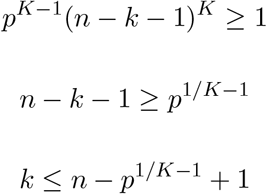

Finally we observe that *p*^1*/K*^ *≤* 1 which allows us to simplify to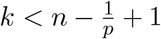. We observe that the strict upper bound for the number of possible simultaneously stored sequences is *n*, achieved only when *p* = 1 (i.e., a complete graph)–in this case, every neuron in each ensemble can be used in a distinct sequence. We also observe that as the network becomes more sparsely connected (i.e., *p→*0), fewer sequences can be stored simultaneously. Finally, for “typical” values of network parameters *N, K* we have *k ≪ K*!, indicating only a small subset of possible sequences in the network can be stored simultaneously.

If redundancy in sequence representations is required (i.e., each sequence must be represented by *t* chains), then divide the RHS by *t*

### Average case performance

We can extend the above argument to the average case, taking the assumption that chains are chosen uniformly at random.

Fix a first chain through the network. Then the probability a second chain uses none of the neurons of the first is

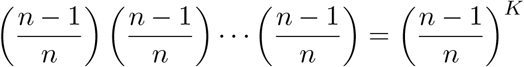

The probability that the *k*-th chain uses none of the neurons of the first *k* − 1 is similarly

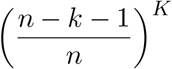

This is a variant of the Birthday Problem, with probability

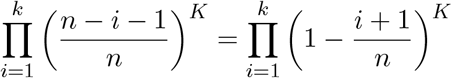

We can use the approximation *e*^−*x*^ *≈* 1 − *x* for *x ≪* 1

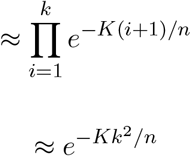

Setting this expression equal to 0.5, we get

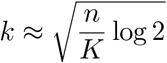

or 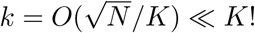. We remark that the best case storage capacity scales linearly with the representational redundancy *n*, while the average case storage capacity scales only with 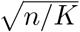. Moreover, the quantity has implicit dependence on *p* in that it only holds when sequences are realizable, which we calculated in Theorem 4 is true w.h.p. when *N > Kp*^1*/K*−1^

